# Comparative metabolism of the *Alternaria* toxins altenuene and tentoxin in rat and human primary hepatocytes

**DOI:** 10.64898/2026.05.11.724251

**Authors:** Eszter Borsos, Charlène Gendre, Mariam Mahdjoub, Elisabeth Varga, Estelle Dubreil, Jérôme Henri, Ludovic Le Hegarat, Doris Marko

## Abstract

The ubiquitously occurring food contaminants altenuene (ALT) and tentoxin (TEN) are recognized as emerging *Alternaria* mycotoxins, yet substantial data gaps remain when it comes to their toxicological behavior and toxicokinetic characteristics. This study aimed to compare and generate quantitative data on their hepatic metabolism and to obtain semi-quantitative insights into their metabolite profiles. To this end, primary rat and human hepatocytes were incubated with 10 µM ALT or TEN over multiple time points up to 4 h. Both substrate depletion and metabolite identification revealed pronounced interspecies differences. The extent of ALT metabolism was significant, with an 88% and 57% decrease in rat and human hepatocytes after 4 h, respectively. In contrast, TEN showed extensive biotransformation in rats (67%) but only modest turnover in humans (27%) over the same period. Hepatocellular clearances were consistently higher for ALT than TEN, with hepatic extraction ratios indicating intermediate extraction for ALT and low extraction for TEN. High-resolution mass spectrometry combined with targeted analysis of selected metabolites annotated phase II conjugation as the predominant metabolic pathway for ALT and phase I oxidative metabolism for TEN, including mono- and double-metabolized species for the latter. Overall, these results provide a comprehensive characterization of ALT- and TEN-metabolism in hepatocytes, offering a foundation for future studies on their toxicological relevance and impact on human health.

## 1. Introduction

Fungi of the genus *Alternaria* are frequent contaminants of plant-based food commodities, particularly cereals, oilseeds, and fruits (Ostry 2008). Among the secondary metabolites they produce, over 70 are known to elucidate adverse effects on vertebrates and are therefore known as *Alternaria* toxins. These substances show a wide variety in structure and toxicological properties, but fall collectively under the category of emerging mycotoxins, as they are not covered by either regulatory limits or monitoring programs despite their increasingly observed occurrence in foodstuffs (Gruber-Dorninger *et al*. 2017). Although risk assessment has traditionally focused on major *Alternaria* toxins such as alternariol (AOH), additional compounds including the dibenzo-α-pyrone altenuene (ALT) and the cyclic tetrapeptide tentoxin (TEN) are consistently detected in occurrence surveys (EFSA 2011). The European Food Safety Authority (EFSA) has highlighted ALT and TEN as understudied mycotoxins in its assessment of secondary *Alternaria* metabolites, mentioning insufficient information on their toxicokinetics and metabolism (EFSA 2016).

ALT and TEN were detected in various food matrices, including fruits, tomato-based products, cereals, and oilseeds, often co-occurring with other mycotoxins, as reviewed by Zhang *et al*. (2025). While several *in vitro* studies have classified both compounds as non-genotoxic (Louro *et al*. 2024; Schrader *et al*. 2001, 2006), there are indications of cytotoxic or hepatotoxic effects. For example, ALT has demonstrated cytotoxicity in the human colon carcinoma cell line HCT116 captured by the sulforhodamine B assay, with an half maximal inhibitory concentration (IC_50_) value of 3.13 μM (Xiao *et al*. 2014). In turn, TEN has been associated with hepatotoxicity through the downregulation of relevant gene expression in differentiated HepaRG cells, potentially contributing to an increased risk of cholestasis and cirrhosis (Hessel-Pras *et al*. 2019).

Information on the hepatic metabolism of ALT and TEN remains notably limited. Pfeiffer *et al*. (2008) demonstrated that ALT is a substrate for oxidative metabolism by human hepatic cytochrome P450 enzymes, preferably the isoform CYP2C19. Across all investigated isoforms, the major metabolite was tentatively annotated as 8-hydroxy-ALT. A comparable pattern was observable when pig, rat, and human liver microsomes (1 mg/mL protein) were treated with 50 µM ALT for 40 min in the presence of a nicotinamide adenine dinucleotide phosphate (NADPH)-generating system (Pfeiffer et al. 2009a). Furthermore, incubation of precision-cut rat liver slices with 200 µM ALT for 24 h at 37 °C also revealed evidence of oxidative metabolism (11.6%), although glucuronide conjugation (49.8%) represented the predominant biotransformation pathway in this *ex vivo* model (Pfeiffer et al. 2009a).

Early work by Delaforge *et al*. (1997) has described the cytochrome P450-dependent oxidative metabolism of TEN mediated primarily by CYP3A isozymes from rat and human liver. Hydroxylation and *N*-demethylation were observed as the major metabolic pathways. A complementary study by Perrin *et al*. (2011) confirmed the formation of the same metabolites in hepatic microsomes from six mammalian species (rat, mice, rabbit, cow, sheep, and human). Furthermore, they demonstrated that oxidation occurs selectively at the *N*-methyl-alanyl residue, forming a transient *N*-hydroxymethyl intermediate that spontaneously undergoes deformylation to yield the *N*-demethylated product (deMe-TEN).

To date, no comprehensive study exists on the hepatic metabolism of ALT and TEN in cellular model systems, which would provide realistic insight into the metabolic profile of these *Alternaria* mycotoxins following the interplay of phase I and phase II biotransformation reactions. Moreover, a substantial data gap remains regarding the kinetic aspects of their metabolism. This data would be essential to understand metabolic activation and/or detoxification pathways, which in turn inform toxicokinetics, biomonitoring, and hazard assessment. To address this gap, we investigated the time-dependent metabolism of ALT and TEN in primary rat (PRH) and human (PHH) hepatocytes over multiple incubation times (0–4 h), using liquid chromatography – tandem mass spectrometry (LC-MS/MS) for quantification and ultra-high performance liquid chromatography – high resolution mass spectrometry (UHPLC-HRMS) for the structural annotation of resulting metabolites. These data provide novel insights into their biotransformation and establish the foundation for subsequent toxicological evaluation and regulatory risk assessment.

## 2. Materials and methods

### 2.1 Chemicals and biological materials

#### 2.1.2. Chemicals and reagents

Altenuene (batch number 0701458-3) and tentoxin (batch number 070146-1) were purchased from Cayman Chemical (Michigan, USA) with purities greater than 98% and 99%, respectively. Dimethyl sulfoxide (DMSO) was purchased from Sigma (St. Quentin-Fallavier, France). Stock solutions at 20 mM were prepared in DMSO.

For sample preparation and subsequent LC-MS/MS analysis, CHROMASOLV LC−MS-grade acetonitrile (ACN) and methanol (MeOH) from Honey-well Riedel-de Haen (Seelze, Germany), as well as LiChrosolv Hypergrade MeOH from Supelco (Munich, Germany) were used. HiPerSolv CHROMANORM and OPTIMA LC-MS grade water were purchased from VWR International (Radnor, PA, USA) and Thermo Fisher Scientific (Waltham, MA, USA), respectively. As an acidic eluent modifier, ROTIPURAN LC−MS-grade formic acid (Carl Roth, Karlsruhe, Germany) was used. Certified reference solutions of ALT and TEN from the brand Biopure were purchased from Romer Labs (Tulln, Austria).

#### 2.1.3 Primary hepatocyte source and required media with supplements

Cryopreserved (Sprague-Dawley) rat primary hepatocytes (grade S for suspension assays, batch HEP134067) and cryopreserved human primary hepatocytes (10 Caucasian donor mixed gender pooled: 5 men and 5 women) (grade S for suspension assays, batch HEP190023-TA05) were purchased from Wepredic (Biopredic International, Saint-Grégoire, France). Thawing medium (MIL130), treatment medium composed of basal hepatic cell medium (MIL600) supplemented with an aliquot of additives (ADD222) composed of penicillin 100 UI/mL, streptomycin 100 µg/mL, bovine insulin 4 µg/mL and hydrocortisol 50 µM (ADD222) were purchased from Wepredic (Biopredic International, Saint-Grégoire, France).

### 2.2 Treatment of primary hepatocytes with ALT and TEN

#### 2.2.1 ALT and TEN kinetics with PRH and PHH

The cryopreserved vials of hepatocytes were thawed in a water bath at 37 °C and transferred to a thawing medium heated to 37 °C (MIL130), according to supplier recommendations. After centrifugation at 100 *g* and 180 *g* for rat and human hepatocytes, respectively, the supernatant was removed, and the cells were re-suspended in 3 mL of treatment medium. Viability after reconstitution was determined using Trypan blue at 0.4% and was always above 80%. Then, in a 5-mL glass tube, hepatocytes were incubated in suspension, at a final density of 5.10^5^ cells/mL and a volume of incubation of 200 µL, for independent kinetics with ALT and TEN at 10 µM with a final concentration of 0.05% DMSO, at 37 °C with 5% CO_2_ on a microplate shaker. For each time point, T0, 0.5, 1, 1.5, 2 and 4 hours, metabolism was stopped by adding ice-cold (-20 °C) ACN (1:1, *v/v*) and the extraction protocol was conducted as described below. At the end of the incubation period, cell viability was assessed by using the CellTiter-Glo^®^ Luminescent Cell Viability Assay (Promega Corporation, Madison, WI, USA), that measure ATP content. For each time point, three technical hepatocyte replicates were performed.

#### 2.2.2 Extraction of ALT and TEN

Once metabolism was stopped by ACN, the tubes were sonicated for 10 min in a water bath, then the suspensions were transferred to 1.5 mL low-binding Eppendorf tubes and centrifuged for 5 min at 10,000 *g* at room temperature. The supernatants were filtered through 0.22 µm polyvinylidene difluoride (PVDF) filter (Sigma-Aldrich, Ireland) and transferred to 2 mL amber glass vials. The samples were stored at -80 °C until analysis.

#### 2.2.3 Stability assays for ALT and TEN

The stability of ALT and TEN were determined by using 5 mL glass tubes containing William’s E medium during T0, 4 and 75 hours of incubation at 10 µM. After each incubation period, metabolism was stopped by adding ice-cold (-20 °C) ACN (1:1, *v/v*) and the extraction protocol was conducted. For each time point, three technical hepatocyte replicates were performed. The samples were injected into the UHPLC-HRMS Q-Exactive Plus spectrometer (Thermo Scientific) according to the UHPLC-HRMS method described below in 2.4.2. Ratio of area (ALT neg: *m/z* 291.08741; TEN neg: *m/z* 413.21943) between different incubation periods and T0 were reported to evaluate the stability of ALT and TEN in William’s E medium incubated at 37 °C.

### 2.3 Parent toxin decrease in ALT or TEN-treated rat and human hepatocytes

#### 2.3.1. Sample preparation for the targeted analysis

Samples of three independent biological hepatocyte experiments were obtained for the targeted analysis as described in subsection 2.2.1./2. Subsequently, they were further diluted with MeOH-H_2_O (1:9, *v/v*; 1:5 and 1:10 for ALT and TEN, respectively) in analytical triplicates to ensure that the measured analytes concentrations fall within the calibration range of ALT or TEN.

#### 2.3.2. Quantification of ALT and TEN with high-performance liquid chromatography coupled to tandem mass spectrometry (LC-MS/MS)

The targeted measurements of ALT and TEN were conducted on an ultra-high-performance liquid chromatography instrument (1290 Infinity II of Agilent Technologies, Santa Clara, CA, USA) coupled to a QTRAP 6500+ mass spectrometer (SCIEX, Framingham, MA, USA) with an electrospray ionization (ESI) interface.

Chromatographic separation was achieved on an Ascentis^®^ Express C18 column (100 mm × 2.1 mm, 2.7 µm; Supelco, Munich, Germany) equipped with a C18 pre-column (SecurityGuard, 4.0 mm × 2.0 mm; Phenomenex Ltd., Aschaffenburg, Germany). A multistep gradient elution was applied at a flow rate of 0.4 mL/min, with an aqueous and methanolic formic acid solution (both 0.1%) as eluent A and B, respectively. During the first minute, the column was rinsed with 10% eluent B. Subsequently, the content of eluent B was linearly raised to 80% until the seventh minute, and then steeply increased to 100% within the next half a minute. Thereafter, the column was washed with 100% eluent B for 1.5 min. Lastly, the column was re-equilibrated in the initial conditions, reaching an overall run time of 12 min. The autosampler compartment was cooled to 10 °C, while the column oven temperature was maintained at 30 °C.

The tandem mass spectrometer was operated in multiple reaction monitoring mode under negative ionization mode, allowing the detection of analytes in their deprotonated forms. External calibration using solvent-based standards was applied for the quantification of the parent toxins with a linear range of 10-1000 µg/L and 3-100 µg/L for ALT and TEN, respectively. Data acquisition was performed using the Analyst (version 1.7., Sierra Analytics, CA, USA) software or Sciex OS (version 3.3.1.43) and data evaluation was carried out with the open-source software Skyline (version 25.1., MacCoss Lab, University of Washington, WA, USA) or with Sciex OS.

#### 2.3.3. Evaluation of the method performance

A solvent-based spiking solution was prepared for ALT and TEN and used to spike samples before and after sample preparation to assess recovery and matrix effects. For each condition, three independent sample sets were produced and analyzed.

Extraction recoveries (RE) were calculated as the ratio of the average ALT or TEN concentrations measured in samples spiked prior to extraction with those spiked after extraction. Particular focus was laid on potential losses during syringe filtration, as *Alternaria* mycotoxins are known to adsorb to various plastic materials, including polyvinylidene difluoride (PVDF) (Aichinger *et al*. 2020).

Potential signal suppression or enhancement (SSE) caused by the cell culture medium was evaluated by dividing the concentration of ALT or TEN in post-spiked medium by the concentration measured in a solvent-based solution of the same spiking level. As no quantifiable ALT or TEN was detected in the matrix blanks (pure medium), no background subtraction was required for RE and SSE calculations. Finally, the toxin-containing media used for hepatocyte treatment – both with and without syringe filtration – were analyzed to determine the actual initial toxin concentrations.

#### 2.3.4. Calculation of the in vitro and in vivo intrinsic clearance values

Based on the decrease of ALT and TEN observed in primary rat and human hepatocytes, the metabolic stability of these compounds was assessed. In this approach, the nominal incubation concentration (10 μM) was selected sufficiently low to fall below the expected Michaelis constant (K_m_), ensuring linear substrate-depletion kinetics. At the same time, a slightly higher concentration was chosen compared with other studies (0.1-1 µM) to enable the detection of metabolization products. The first-order elimination rate constant (k_inc_, [1/h]) was obtained from the negative slope of the log-linear plot of substrate concentration (ln(c_t_/c_0_)) versus time, following linear regression (Baranczewski *et al*. 2006).

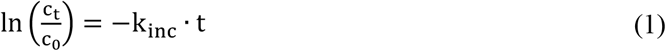

where c_t_is the ALT/TEN concentration [µM] at time t[h] and c_0_is the initial concentration [µM].

Subsequent calculations, including *in vitro* intrinsic clearance, whole-liver intrinsic clearance, and hepatic clearance (Equations (2)-(4)), were performed according to Sodhi and Benet (2021). The *in vitro* intrinsic clearance (CL_int, *in vitro*_, [mL/(h·10^6^ cells)]) was calculated from k_inc_ using equation (2), incorporating the incubation volume (V_inc_,[mL])and the number of hepatocytes in the *in vitro* incubation (N_cells, *in vitro*_, [10^6^ cells]).

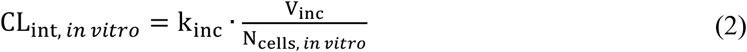

The intrinsic clearance for the whole liver (CL_int, *in vivo*_, [mL/(h·kg bw)]) was obtained by scaling *in vitro* clearance with the *in vivo* hepatocyte content (N_cells, *in vivo*_), as described in equation (3).

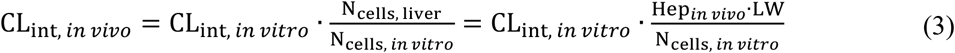

where Hep_*in vivo*_ stands for the *in vivo* hepatocellularity [cells/mL] and LW describes the relative liver weight in g/kg body weight (bw). N_cells, *in vitro*_ was set to 10^6^ cells, consistent with the normalization of CL_int, *in vitro*_.

Hepatic clearance (CL_H_, [L/(h·kg bw)]) was calculated based on the well-stirred model, assuming the fraction unbound in plasma (f_u,B_) to be 1 (equation (4)).

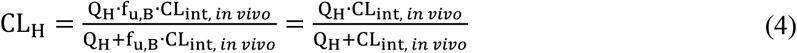

where Q_H_ represents the hepatic blood flow [L/h·kg bw)].

Finally, the hepatic extraction ratio (E_H_) was calculated based on equation (5).

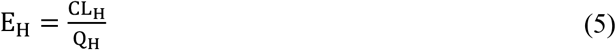

### 2.4 Hepatic metabolites of ALT and TEN

#### 2.4.1. In-silico prediction of possibly occurring metabolites

ADMETpredictor^®^13.0 (AP) is an *in silico*-based prediction module using structure-property relationships to predict a wide range of ADMET (absorption, distribution, metabolism, excretion, and toxicity) properties. Within its 9 modules, AP can predict phase I and phase II metabolism for a given molecule. The 2D structure, in SDF format, was retrieved from PubChem and loaded into AP. For the submitted chemical structure, AP predicted primary and secondary metabolites, suggesting potential metabolic pathways and a comprehensive aspect of the metabolic fate of the compound.

#### 2.4.2 Metabolic profiling using UHPLC-HRMS analysis

A metabolic profiling method was applied in order to detect potential mono- and double-conjugated metabolites in primary hepatocytes for one biological hepatocyte replicate per time point (0 h, 0.5 h, 1 h, 1.5 h, 2 h, 4 h for PRH and 0 h, 1 h, 4 h for PHH) without any additional dilutions. The samples were all analyzed using a Vanquish Flex binary UHPLC system (Thermo Fisher Scientific, Waltham, MA, USA) with an Acquity UPLC HSS T3 column (150 mm × 1 mm, 1.8 µm, 100 Å). The column temperature was maintained at 35 °C, the flow rate was at 0.080 mL/min and the injection volume was 5 µL. Mobile phases were composed of LC-MS grade water containing 0.1% formic acid (eluent A) and acetonitrile/methanol (80:20, *v/v*) containing 0.1% formic acid (eluent B). The gradient elution conditions were set as follows: 0-2.5 min linearly increased from 2% to 10% B; 2.5-9 min linearly increased to 45% B; 9-19 min linearly increased to 100% B and held for 6 min; 25-26 min linearly decreased to 2% B and kept up to 32 min to allow the system to equilibrate. The UHPLC was coupled to a Q-Exactive Plus mass spectrometer (Thermo Fisher Scientific, Waltham, MA, USA). Full MS scan data in both electrospray positive and negative modes (ESI+, ESI-) were acquired at a resolving power of 70,000 full width at half maximum (FWHM) at *m/z* 200 with a mass range of *m/z* 100–1000. The automatic gain control (AGC) target was set at 3×10^6^ ions, with a maximum injection time of 100 msec.

A heated electrospray ionization source H-ESI II probe was used with parameters set as follows: sheath gas (nitrogen) flow rate 45 arbitrary units; auxiliary gas (nitrogen) flow rate 10 arbitrary units; sweep gas (nitrogen) flow rate 10 arbitrary units; spray voltage 3.50 kV; capillary temperature 320 °C. MS/MS data were acquired in the Orbitrap cell in DDA (Data Dependent Acquisition) mode. Inclusion list (Table S1) containing the exact masses of the parent compounds was used to obtain product ion spectra with a mass resolution set at 17,500 FWHM (*m/z* 200). MS/MS data fragments were obtained by higher-energy collisional dissociation (HCD) activation with a collisional energy of 20, 40 and 70% and an isolation width of *m/z* 1.5. The dd-MS2 parameters were set as follows: the 5 most intense precursor ions were fragmented; with a dynamic exclusion of 30 sec; AGC targets were set to 1×10^5^; maximum ion accumulation times were set to 64 msec; the data were acquired in profile mode. External mass calibrations were performed weekly. The acquisition software was Xcalibur version 4.4. Data processing was performed using the TraceFinder EFS software version 5.1 for peak-detection algorithm by setting target mass tolerance at ± 5.0 ppm.

#### 2.4.3. Targeted analysis of selected ALT and TEN metabolites via LC-MS/MS

The list of ALT and TEN metabolites of interest for the targeted method was compiled based on available literature, *in silico* metabolite prediction, as well as HRMS analysis of the incubation samples described above. Three biological hepatocyte replicates per time point (0 h, 0.5 h, 1 h, 2 h and 4 h for PRH and PHH and additionally 1.5 h in case of PRH) without any additional dilutions were used for these investigations. In the absence of reference standards for the biotransformation products of ALT and TEN, method optimization relied on the repeated injection of selected incubation samples containing the peaks of interest. The samples were injected *via* LC autosampler, underwent chromatographic separation, and critical MS parameters, including declustering potential (DP) and collision energy (CE), were manually tuned to enhance signal intensity. The final transition list, including the optimized parameters, can be found in Table S2.

## 3. Results and Discussion

### 3.1 Stability of ALT and TEN

The stability results for ALT and TEN toxins after 1 and 75 hours of incubation at 35 °C are provided in the Supplementary materials (Figure S1). After 75 hours of incubation, both toxins were stable with no significant decrease compared to T0 and the treatment solution.

### 3.2 Treatment of primary hepatocytes with ALT and TEN

#### 3.2.1 Evaluation of the method performance

The extraction recovery (RE) of the sample preparation process showed mean values of 103–107% for ALT and 98–101% for TEN (Figure S2). These results were deemed acceptable, as the deviations from 100% were comparable to the variability observed between replicate measurements under identical conditions. Thus, the differences attributable to recovery did not exceed normal method repeatability. Potential analyte losses due to syringe filtration were also assessed, given that both ALT and TEN is known to adsorb to PVDF filters (Aichinger *et al*. 2020). However, no significant decrease of either analyte was detected after filtration, suggesting that this sample preparation step does not compromise quantification, most likely due to the minimal contact time with the filter material.

Analyte-dependent matrix effects were observed, indicating no considerable effects for TEN and slight signal suppression for ALT (Figure S2). The latter is consistent with the lower dilution factor and consequently higher matrix proportion used for ALT, given that its calibration range spans concentrations an order of magnitude higher than those of TEN. Nevertheless, the observed suppression (10%) falls within ranges commonly considered acceptable for quantitative LC–MS/MS analyses (European Medicines Agency 2023).

#### 3.2.2 Parent toxin decrease and hepatic intrinsic clearance

No significant decrease in cell viability compared to DMSO (0.05%) was observed in PRH and PHH after 4 hours of incubation with ALT and TEN (Figure S3).

In rat hepatocytes, the initial ALT concentration (Figure S4) was almost entirely depleted after 4 h of incubation at 10 µM, showing an 88% reduction (Figure S5). In contrast, the depletion of ALT was slower in human hepatocytes, with only a 57% decrease in the parent compound over the same period.

In rat hepatocytes, TEN was extensively biotransformed after 4 h of incubation at 10 µM, with 67% decrease, whereas only a small fraction of TEN was depleted in human hepatocytes (27% decrease). Notably, in humans, the measured TEN concentrations exhibited a non-monotonic decrease over time, in contrast to the consistently declining trends observed in the other three cases. This irregularity is potentially attributable to the relatively small overall decrease in TEN concentrations in humans, where experimental variability – such as pipetting or sampling inaccuracies – may have had a proportionally greater impact on the measurement outcome.

Across species, ALT displayed markedly higher hepatic clearance than TEN (Table 1). The corresponding hepatic extraction ratios (E_H_) for ALT – 55% and 47% in rats and humans, respectively – categorize it as a compound with intermediate metabolism. TEN, by contrast, exhibited lower hepatic clearance across species, with hepatic extraction ratios of 39% in rat and 26% in human primary hepatocytes. These values designate TEN as a slowly cleared toxin in the human liver. The classification framework stated above is based on commonly applied thresholds, defining compounds with high (E_H_ > 0.7), intermediate (0.3 < E_H_ < 0.7), and low (E_H_ < 0.3) hepatic extraction (Hebert 2013).

**Table 1.** *In vitro* to *in vivo* extrapolation (IVIVE) of the obtained kinetic data for altenuene (ALT) and tentoxin (TEN) and applied parameters.

| Parameter | Unit | Short | ALT |  | TEN |  |
| --- | --- | --- | --- | --- | --- | --- |
|  |  |  | Rat | Human | Rat | Human |
| *In vitro* hepatocellularity | cells/mL | Hep<sub>*in vitro*</sub> | $5 \cdot 10^5$ | $5 \cdot 10^5$ | $5 \cdot 10^5$ | $5 \cdot 10^5$ |
| Incubation volume | mL | V<sub>inc</sub> | 0.2 | 0.2 | 0.2 | 0.2 |
| *In vitro* elimination rate constant | 1/h | k<sub>inc</sub> | 0.54 | 0.21 | 0.28 | 0.08 |
| *In vivo* hepatocellularity | cells/g liver | Hep<sub>*in vivo*</sub> | $1.17 \cdot 10^8$ | $1.39 \cdot 10^8$ | $1.17 \cdot 10^8$ | $1.39 \cdot 10^8$ |
| Relative liver weight | g/kg bw | LW | 40 | 21 | 40 | 21 |
| Liver blood flow | L/(h·kg bw) | Q<sub>H</sub> | 4.2 | 1.4 | 4.2 | 1.4 |
| *In vitro* intrinsic clearance | mL/(h·10<sup>6</sup> cells) | CL<sub>int, *in vitro*</sub> | 1.1 | 0.4 | 0.6 | 0.2 |
| Scaled intrinsic clearance | mL/(h·kg bw) | CL<sub>int, *in vivo*</sub> | 5060 | 1254 | 2662 | 486.9 |
|  | L/(h·kg bw) |  | 5.06 | 1.25 | 2.66 | 0.49 |
| Extrapolated *in vivo* hepatic clearance | L/(h·kg bw) | CL<sub>H</sub> | 2.30 | 0.66 | 1.63 | 0.36 |
| Hepatic extraction ratio | % | E<sub>H</sub> | 55 | 47 | 39 | 26 |

A minimum substrate depletion of 10-20% is widely accepted as the threshold for reliable clearance estimation in hepatocytes (Sodhi and Benet 2021), corresponding to an intrinsic clearance cutoff of ∼0.11 mL/(h·10^6^ cells) (CL_H_ ≈ 0.26 L/(h·kg)) in our system. After 4 h of incubation in human hepatocytes, TEN showed ∼27% turnover (Figure S5B), exceeding the depletion and clearance threshold mentioned above and thus supporting the validity of the IVIVE values. However, the depletion–time profile did not show an ideal first-order kinetics across all five time points, and two data points deviated from log-linearity (Figure *2*D). To obtain a more reliable slope estimate, the linear regression was therefore restricted to three time points, which formed the most consistent quasi-log-linear segment.

The intrinsic clearance we observed for TEN in humans did not enter the “ultra-low clearance” domain (CL_int,*in vitro*_ ≤ 2.5 µL/(min·10^6^ cells) corresponding to 0.15 mL/(h·10^6^ cells)) as defined by Di and Obach (2015). To acquire reliably measurable decreases for compounds falling below this threshold typically requires adaptation of the assay setup, including elevated *in vitro* hepatocellularity, lower substrate levels, or extended incubation time. Due to the practical limitations of these solutions, specialized strategies – such as hepatocyte relay assays or long-lived 3D hepatocyte platforms – could be introduced to obtain reliable turnover measurements (Di and Obach 2015). Since TEN remained slightly above this ultra-low-clearance boundary, and to be able to make comparisons between species and between toxins under the same conditions, these measures were not applied in the present study. In contrast to TEN in humans, the other three investigated toxin-species combinations exhibited clearly detectable turnover well above 20%, yielding substantially more robust and higher-confidence clearance estimates.

Despite the structural dissimilarity between ALT and TEN, both toxins were metabolized faster in (Sprague-Dawley) rat than in human hepatocytes. Rat *in vivo* intrinsic clearance values were approximately four-fold higher for ALT (5.1 vs. 1.25 L/(h·kg)) and about five-fold higher for TEN (2.7 vs. 0.5 L/(h·kg)) than that in human hepatocytes. This species divergence underscores the importance of relying on human-relevant systems when extrapolating toxicokinetics for risk assessment. Using rat-derived clearance to infer human metabolism can overestimate human metabolic efficiency and, therefore, underestimate internal exposure and toxicological risk.

Published biotransformation studies provide qualitative context for our findings in primary hepatocytes. For ALT, microsomal incubations demonstrated clear phase I turnover, with Pfeiffer *et al*. (2009) reporting approx. 25% depletion in rat and 8% in human liver microsomes over 40 min at 1 mg/mL protein in the presence of an NADPH-generating system. These interspecies differences agree with the faster ALT metabolism observed in rat hepatocytes in the present study. In addition to oxidative metabolism, ALT has been shown to undergo glucuronidation in precision-cut liver slices (Pfeiffer *et al*. 2009). ALT being a substrate of both phase I and II metabolic enzymes support a higher metabolic susceptibility of ALT than TEN, in line with the clearance hierarchy identified in our hepatocyte system.

TEN has likewise been shown to undergo oxidative metabolism in liver microsomes of different species, predominantly *via* CYP3A isoforms (Delaforge *et al*. 1997; Perrin *et al*. 2011), although quantitative depletion data – particularly interspecies comparisons – could not be found. No intrinsic clearance values have been published for both compounds, and direct quantitative conversion between microsomal depletion and hepatocyte intrinsic clearance is not possible due to differences in enzyme abundance, cofactors, and assay design. Nevertheless, the agreement in relative metabolic trends strengthens confidence in the clearance ranking established in this work.

*In vivo* toxicokinetic data for *Alternaria* toxins that could provide a quantitative frame of reference for the present hepatocyte-based clearance data are scarce. Following oral administration of a complex *Alternaria* extract to Sprague–Dawley rats – containing 39.2 µg/kg b.w. ALT –, ALT was not detected in plasma or urine at 3 or 24 h (Puntscher *et al*. 2019). Only ∼7% of the administered dose was recovered in feces after 24 h, which may indicate in addition to the metabolism, a low excretion and/or incomplete recovery; however, no quantitative *in vivo* clearance values are available.

Given the lack of compound-specific data, structurally related dibenzo-α-pyrones such as AOH and alternariol monomethyl ether (AME) can provide contextual information. In the same rat study, both compounds were predominantly recovered in feces (>89%) and were not detectable in plasma, consistent with low oral bioavailability combined with pronounced presystemic elimination (Louro *et al*. 2024; Puntscher *et al*. 2019). In line with this, studies in pigs demonstrated low oral bioavailability (≈9–15%) but very high systemic clearance (≈13–17 L·h^−1^·kg^−1^) for AOH and AME, indicating rapid hepatic elimination once absorbed (den Hollander *et al*. 2025). Compared to these compounds, the lower fecal recovery of ALT suggests a distinct toxicokinetic behavior.

For TEN, no parent compound was detected in plasma at 3 or 24 h after oral administration in rats (1.2 µg/kg b.w.), while only trace amounts were recovered in urine. In contrast, approximately 45% of the administered dose was recovered in feces, indicating that a substantial fraction remained unabsorbed (Puntscher *et al*. 2019). Together, these findings point to a limited oral bioavailability as the primary determinant of systemic exposure. In this context, the moderate intrinsic hepatic clearance observed in human hepatocytes in the present study suggests slower metabolic turnover compared to other *Alternaria* toxins, but the *in vivo* relevance of this finding is likely limited by absorption constraints. Overall, *Alternaria* toxins exhibit highly compound-specific toxicokinetic behavior, and comparison of hepatocyte-derived intrinsic clearance with *in vivo* data requires careful consideration of absorption and first-pass effects.

### 3.3 Hepatic metabolites of ALT and TEN

#### 3.3.1 ADMET in silico prediction

*In silico* ADMET Prediction™ software was used to predict the metabolites of ALT and TEN. ALT was predicted to generate only two metabolites for the phase II reaction from UGT-mediated reactions (Figure S6). In contrast, no phase II metabolites were predicted for TEN, but four metabolites for the phase I reaction were identified and implicated CYP3A4 enzyme (Figure S7). Table S3 summarizes the molecular weight and proposed structure of the putative phase I and II biotransformation metabolites for ALT and TEN.

#### 3.3.2 UHPLC-HRMS results

In this metabolic profiling experiment, UHPLC-HRMS data were processed to detect putative phase I and phase II metabolites from rat and human primary hepatocytes of ALT and TEN incubations. All samples were analyzed in DDA mode, which led to the triggering of MS2 events for the most intense precursor ions. The metabolites are listed in Supplementary Information, in Table S4 and S5 for ALT, and in Table S6 and S7 for TEN. All the peaks corresponding to metabolites of ALT and TEN discussed in the next paragraphs were not detected in the solvent controls, treatments solutions and T0 samples. Extraction ion chromatograms and fragmentation spectra are listed in the Supplementary Information. These metabolites were subsequently included in the targeted LC-MS/MS method to get semi-quantitative information.

##### ALT detected metabolites

As summarized in Figure 3 of Louro *et al*. (2024), ALT can be subjected to phase I oxidative metabolism, possibly followed by phase II conjugation. Additionally, the three free hydroxy groups can be sites of direct conjugation at positions 2, 3, and 7, without prior functionalization. In the current study of PRH and PHH, only phase-II metabolites, proposed as *O*-sulfate (M1) and *O*-glucuronide (M2) conjugation, of ALT were detected. This is in line with the ADMET prediction. Phase I oxidative metabolites depicted in Sprague-Dawley rat and human liver microsomes could not be confirmed (Pfeiffer *et al*. 2009b). M1 (ALT-S, *m/z* 373.0597 [M+H]^+^, C_15_H_16_O_9_S) was observed at 11.6 min and 11.8 min as probable isomers, with retention times slightly lower than the parent compound (12.2 min) and in line with the higher polarity. They were 79.96 Da heavier than ALT corresponding to the addition of sulfate and MS/MS experiments revealed fragments of *m/z* 257.0807 (similar to ALT and corresponding to the loss of two water molecules from the parent compound), 227.0704, 214.0468, 67.2097. M2 (ALT-GlcA, *m/z* 469.1349 [M+H]^+^, C_21_H_24_O_12_) was detected in total at four retention times (8.1 min, 8.3 min (no DDA), 10.7 min and 11.4 min) probably due to different positions of the glucuronide moiety (*OH*-conjugation). M2 was 176.03 Da heavier than the parent compound corresponding to the addition of glucuronic acid. As main MS/MS fragments the masses of *m/z* 293.1021 (corresponding to the parent compound ALT), 275.0913, 257.0807 (similar fragments as ALT and corresponding to the loss of one and two water molecules from ALT, respectively) and *m/z* 67.2100 were observed.

##### TEN detected metabolites

In contrast to ALT but in line with the ADMET prediction, only phase I metabolites of TEN were detected in PRH and PHH. Two mono-reactions already depicted as hydroxylation and *N*-demethylation of tentoxin (Delaforge *et al*. 1997; Perrin *et al*. 2011), were present in both PRH and PHH. M1 (OH-TEN, *m/z* 431.2296 [M+H]^+^, C_22_H_30_N_4_O_5_), was observed at 11.0 min, 11.7 min and 12.8 min as probable isomers, with retention times slightly lower than the parent TEN (14.0 min) and in line with the slightly higher polarity. M1 was 16 Da heavier than the parent compound (addition of a hydroxy group), and MS/MS fragments of *m/z* 302.1490, 217.0969, 189.1022, 132.0809; 58.0661 were detected similar to TEN and corresponding to peptide fragments. M2 (deMe-TEN, *m/z* 401.2188 [M+H]^+^, C_21_H_28_N_4_O_4_), was detected at 13.1 min and 14 Da lighter than TEN (loss of methyl-group). MS/MS fragments of *m/z* 217.0975; 189.1025; 132.0810; 86.0972; 58.0662 were observed similar to TEN peptide fragments. Besides these two already previously reported metabolites (Delaforge *et al*. 1997; Perrin *et al*. 2011), three double-biotransformed metabolites were observed, combining demethylation(s) and hydroxylation: M3 (di-deMe-TEN, *m/z* 387.2033 [M+H]^+^, C_20_H_26_N_4_O_4_, 12.1 min), M4 (OH)_2_-TEN, *m/z* 447.2248 [M+H]^+^, C_22_H_30_N_4_O_6_, 11.9 min), M5 (OH-deMe-TEN, *m/z* 417.2140 [M+H]^+^, C_21_H_28_N_4_O_5_, 10.6 min). In case of M4 and M5 very broad peak shapes were observed, probably due to non-differentiated isomeric forms. These three metabolites were poorly fragmented and contained some TEN similar fragments as *m/z* 189.1024, 132.0810 and 86.0972, but for full structural elucidation, further experiments out of the scope of the current study would be required.

#### 3.3.3 Targeted analysis of selected ALT and TEN metabolites via LC-MS/MS

In accordance to the *in silico* predictions (Figure S6) and HRMS analysis, the targeted approach found that ALT exclusively undergoes conjugation reactions *via* glucuronidation in both species (Figure 3A-B) and sulfation in PRH (Figure 3A). Whereas the *in silico* software proposed two glucuronide metabolites, three chromatographic peaks were detected in both species, as expected due to the structural features of ALT. The metabolite eluting at the latest retention time (5.5 min) displayed the highest peak area and can therefore be considered the predominant glucuronide metabolite in both species (assuming similar ionization efficacies). Notably, a fourth putative metabolite was observed in the HRMS analysis hinting that the targeted method could not resolve two isomers. This may be explained by differences in the chromatographic method, different columns used and particularly the shorter total run time (12 min vs. 32 min, respectively). However, as ALT only contains three structurally plausible glucuronidation sites and no MS/MS was obtained for all four ALT-GlcA peaks in HRMS, the production of an additional glucuronide cannot be conclusively confirmed. Moreover, due to the lack of available reference materials and spectral libraries, the identity of these metabolites cannot be fully elucidated.

An additional biotransformation pathway observed in PRH was sulfation. Two ALT-S metabolites were found in the LC-MS/MS analysis (Figure 3 and S8), which were not detected in PHH. Although rats, especially male rats, are considered to possess a higher hepatic sulfotransferase (SULT) activity than humans (O′Brien *et al*. 2004), the presence of ALT-S in the metabolic mixture of PHH cannot be excluded. Still, due to the overall lower reaction rate, these presumed metabolites were not detectable.

While the oxidative metabolism of ALT was reported in the literature in NADPH-fortified liver microsomes (Pfeiffer *et al*. 2009) and several human recombinant CYP isoforms – such as CYP2E19 – were capable of mediating the phase I metabolism of ALT (Pfeiffer *et al*. 2008), no hints towards oxidative ALT metabolism were found in this study with the applied analytical conditions. Our finding of glucuronidation representing the main metabolic pathway of ALT in PRH, is in line with the observation of Pfeiffer *et al*. (2009) in precision-cut liver slices, where approx. 50% of the ALT treatment concentration was recovered as a glucuronide metabolite. This liver model is more complex than microsomes supplemented with single cofactors and thus provides a more physiologically relevant representation of hepatic metabolism, closely resembling primary hepatocytes and *in vivo* liver function (Guillouzo and Guguen-Guillouzo 2008).

In contrast to ALT, TEN underwent exclusively phase I metabolism in both PRH and PHH, notably demethylation and hydroxylation (Figure 3 and S8). In addition to three hydroxylated metabolites predicted *in silico* and confirmed by HRMS, a fourth one was identified *via* targeted analysis. Besides the primary and secondary carbon atoms of L-leucine (Figure 1, amino acid 2) and the methyl group of *N*-methyl-L-alanine (amino acid 1) predicted by *in silico* modeling (Table S6 and Figure S7), additional hydroxylation sites are present in TEN, most notably the aromatic ring of *N*-methyl-trans-dehydro-phenylalanine (amino acid 3). This structural feature provides a plausible explanation of a further OH-TEN constitutional isomer. Furthermore, demethylated TEN was detected, in accordance with the ADMET and HRMS dataset.

**Figure 1.**
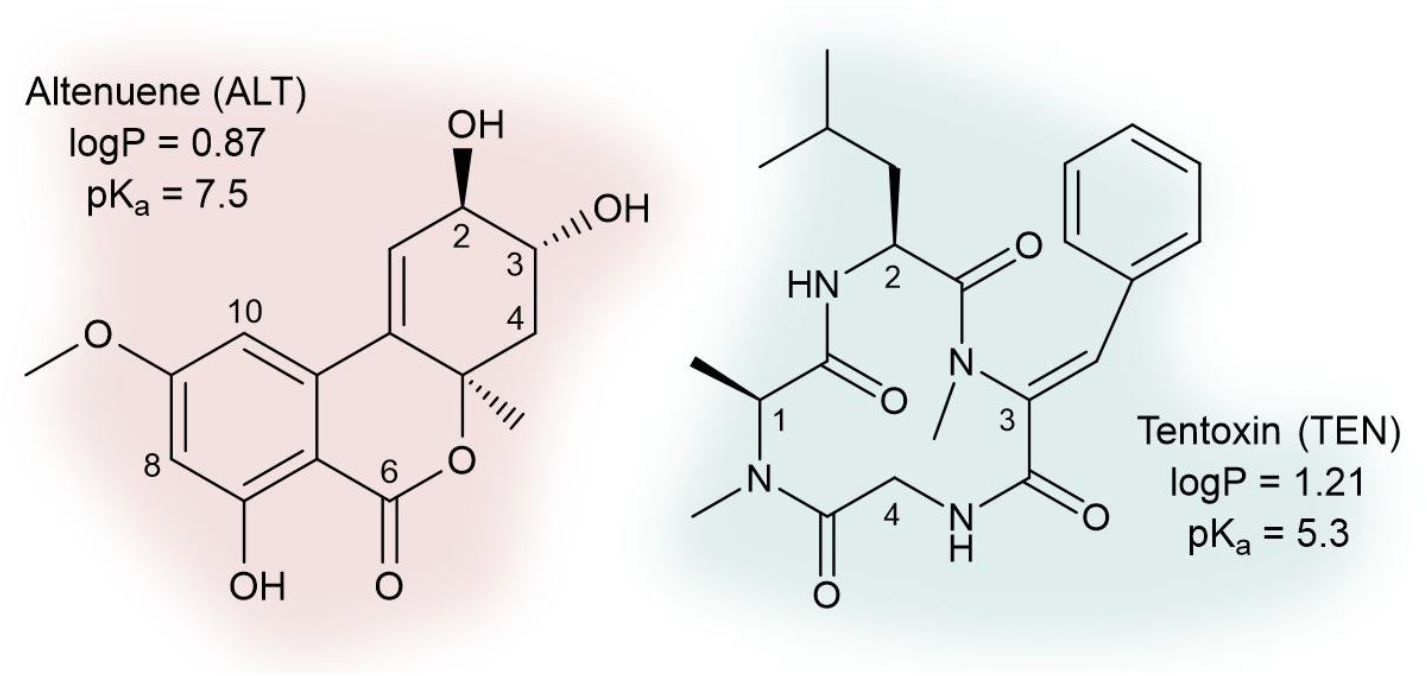
Structure and physical-chemical properties of ALT and TEN. Structures drawn with the software ChemDraw (Revvity Signals Software, Inc. Waltham, MA, USA; v. 23.1.2.7.) and parameters were taken from Tölgyesi *et al*. (2015).

**Figure 2.**
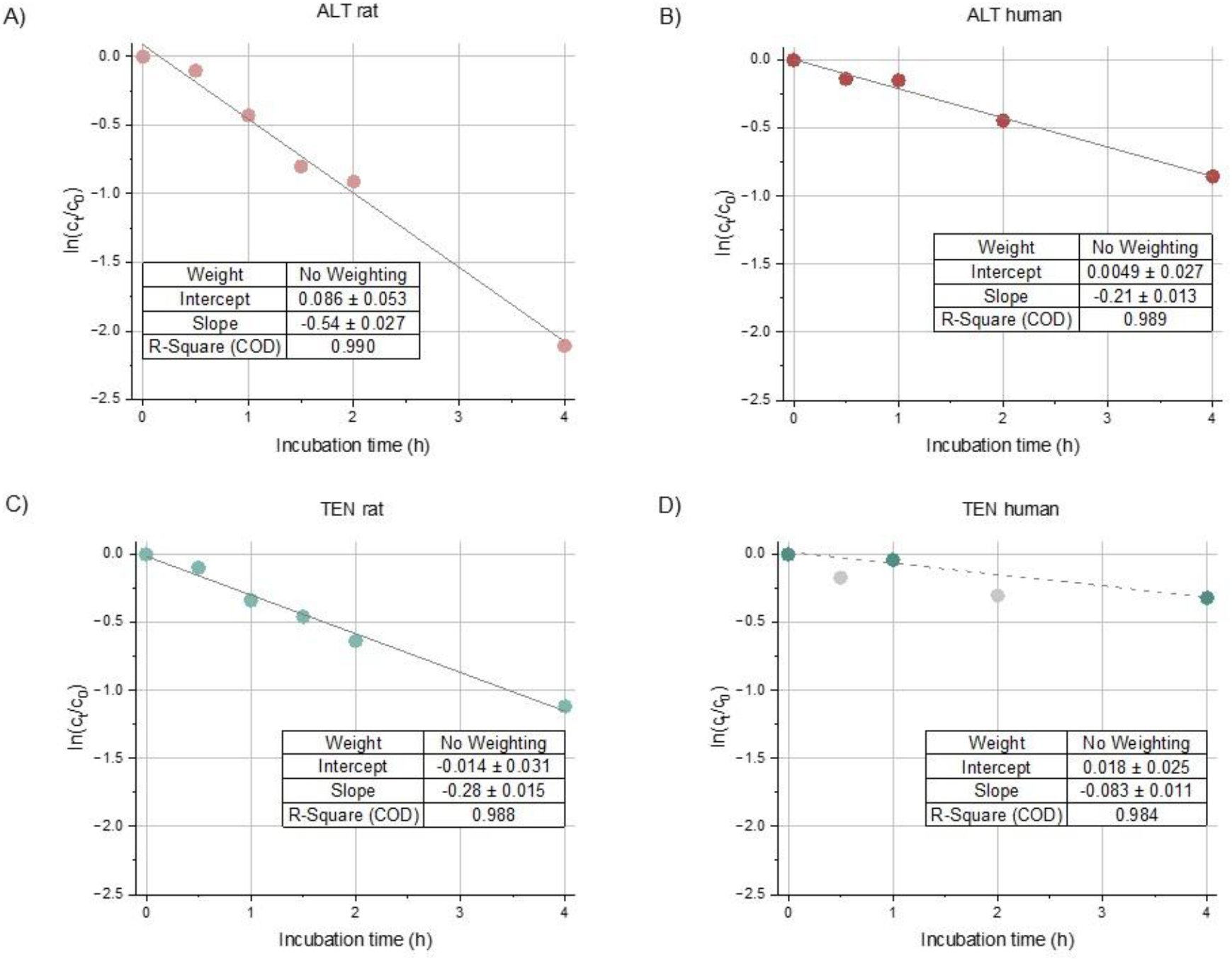
Substrate depletion of ALT (A–B) and TEN (C–D) in rat and human hepatocytes. Regression parameters (slope, intercept, R^2^) are shown in each panel. Each data point represents the mean of three technical analytical replicates from three biological hepatocyte replicates. For human hepatocytes, *c*_0_ was determined from a single biological hepatocyte replicate. Outliers were identified and excluded using Nalimov’s test.

**Figure 3.**
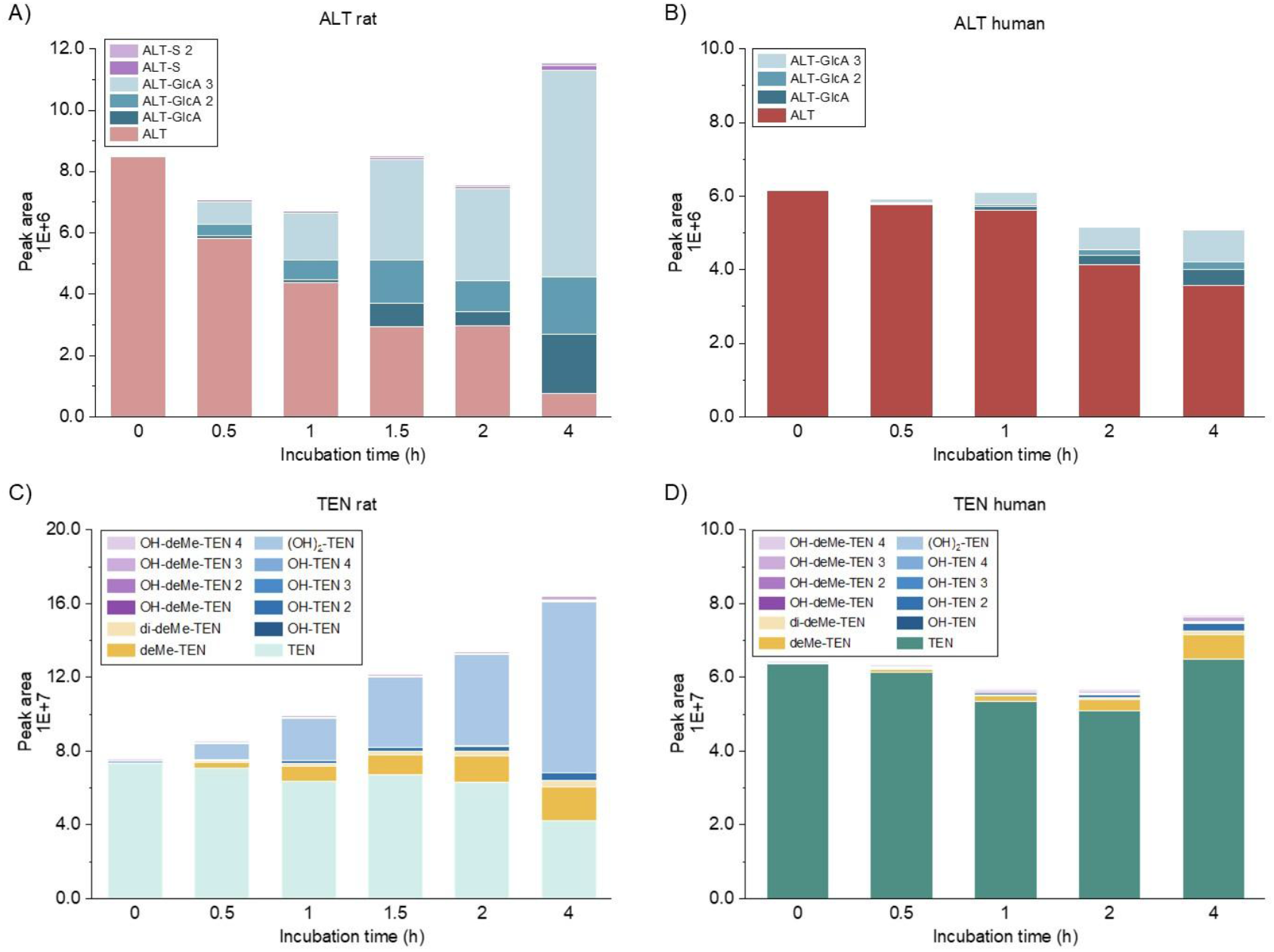
Interspecies differences in the composition of the metabolic mixture produced in primary rat and human hepatocytes after up to 4 h of incubation with 10 μM ALT (A-B) or TEN (C-D) measured *via* LC-MS/MS analysis. Each section shows the average of three independent experiments. Please note the differences in the y-axes.

Pronounced interspecies differences occurred in the relative abundance of individual metabolites. While the dihydroxylated TEN metabolite consistently exhibited the highest peak area in PRH across all time points, demethylated TEN was predominant in PHH, accompanied by an overall more evenly distributed metabolic profile (Figure S8). These findings suggest species-specific preferences in oxidative enzyme activity, which might gain relevance for interspecies extrapolation of TEN toxicity.

Notably, metabolites resulting from multiple subsequent biotransformation steps were also produced, such as hydroxylated-demethylated, as well as double-hydroxylated and double-demethylated TEN derivatives (Figure S8). Although the oxidative metabolism of TEN has previously been reported in liver microsomes from different species, leading primarily to the formation of OH-TEN and deMe-TEN (Delaforge *et al*. 1997; Perrin *et al*. 2011) the present study expands the known metabolic spectrum of TEN by demonstrating the formation of multiple secondary and double-biotransformed metabolites. To our knowledge, this level of metabolic complexity has not been described for TEN before.

As no reference standards are available for any ALT and TEN metabolites, the reported semi-quantitative assessment of metabolite abundance based on peak areas relies on the assumption of comparable ionization efficiencies among structurally related ALT- and TEN-derived analytes, measured under identical matrix and instrumental conditions. However, differences in ionization efficiency may span several orders of magnitude, representing an inherent limitation of the analytical approach (Liigand *et al*. 2018). Despite this limitation, the present data provides some of the first insights into the metabolism of these comparatively understudied mycotoxins in a physiologically relevant hepatocyte model, in which the coexistence of phase I and phase II enzymes is considered. The resulting semi-quantitative metabolic profiles offer a basis for prioritizing metabolite candidates for subsequent toxicological characterization.

## 4. Conclusion

The current study offers a qualitative and quantitative characterization of the hepatic metabolism of the understudied *Alternaria* toxins ALT and TEN in primary rat and human hepatocytes. We observed clear interspecies differences, not only in metabolic rate but also of the resulting metabolite profiles.

Due to their fundamental structural differences, ALT and TEN exhibited distinct metabolic behaviors in primary hepatocytes. ALT showed intermediate intrinsic and hepatic clearance with moderate extraction, while TEN demonstrated low clearance and low extraction in human hepatocytes. Rat hepatocytes metabolized both toxins faster than human hepatocytes, reflecting most common interspecies differences in xenobiotic metabolism.

Semi-quantitative investigations of the occurring metabolites revealed that ALT primarily undergoes phase II conjugation, while TEN was more prone to oxidative phase I metabolism. These findings address a crucial data gap in the toxicokinetics of these poorly studied *Alternaria* toxins and provide quantitative input parameters suitable for physiologically based kinetic (PBK) modeling and potentially supporting future risk assessment efforts.

## Associated content

## Supporting information

The Supporting Information is available free of charge at “xxx to be inserted at a later stage xxx”.

### Author Contributions

All authors have approved the final version of the manuscript. CRediT:

**Eszter Borsos** conceptualization, data curation, formal analysis, investigation, validation, visualization, writing – original draft, writing – review & editing

**Estelle Dubreil** conceptualization, data curation, formal analysis, investigation, validation, visualization, methodology, conceptualization, supervision, project administration, writing – original draft, writing – review & editing

**Charlène Gendre** conceptualization, data curation, formal analysis, investigation, validation, visualization, writing – original draft, writing – review & editing

**Mariam Mahdjoub** formal analysis, writing – original draft, writing – review & editing

**Elisabeth Varga** methodology, conceptualization, supervision, project administration, writing – review & editing

**Jérôme Henri** methodology, conceptualization, supervision, project administration, writing – review & editing

**Ludovic Le Hegarat** methodology, conceptualization, supervision, project administration, writing – review & editing

**Doris Marko** resources, funding acquisition, conceptualization, supervision, writing-review & editing

### Funding

The European Partnership for the Assessment of Risks from Chemicals has received funding from the European Union’s Horizon Europe research and innovation program under Grant Agreement No 101057014 and has received co-funding of the authors’ institutions. Views and opinions expressed are, however, those of the author(s) only and do not necessarily reflect those of the European Union or the Health and Digital Executive Agency. Neither the European Union nor the granting authority can be held responsible for them. Open access funding was provided by the University of Veterinary Medicine, Vienna.

### Notes

The authors declare no competing financial interest.

## Acknowledgements

This research was supported using resources of the VetCore Facility (Mass Spectrometry) of the University of Veterinary Medicine Vienna.

