## Supplementary Material for "Comparative metabolism of the *Alternaria* toxins altenuene and tentoxin in rat and human primary hepatocytes"

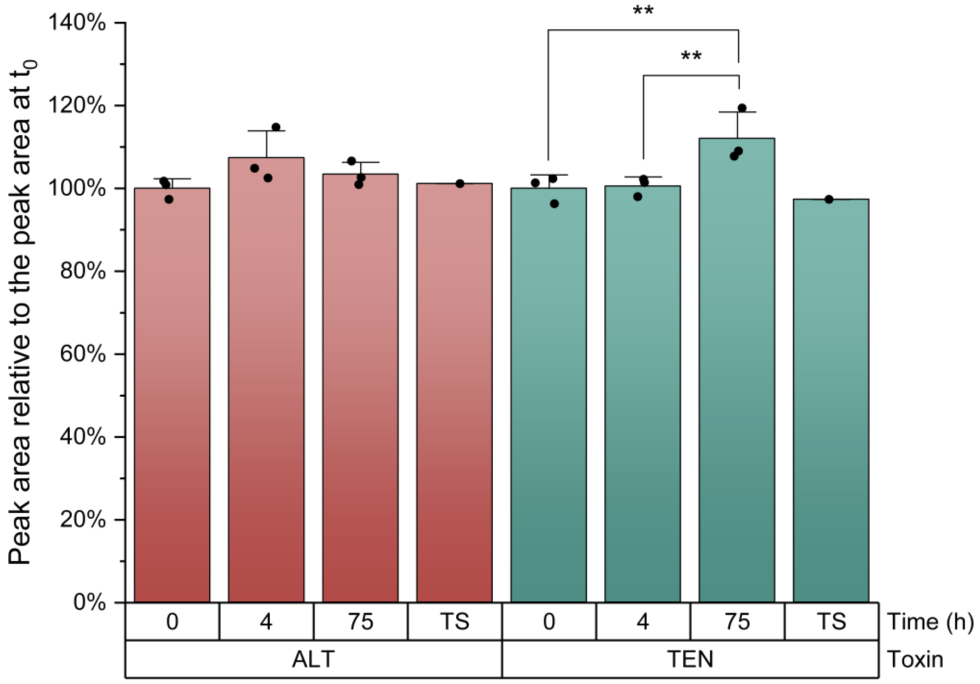


**Figure S1:** Stability of ALT and TEN at 37 °C in William's E medium relative to time point zero ($t_0$). The LC-HRMS method was used in negative mode for this evaluation. Each bar represents the mean value and standard deviation of three replicates which are also depicted. The abbreviation TS stands for the treatment solution (10 µM ALT/TEN), which was only measured in one replicate. After testing for normality according to Shapiro-Wilk, one-way ANOVA, followed by Fisher's LSD post-hoc test was performed. The significance levels are marked as follows: * → $0.01 < p < 0.05$; ** → $0.001 < p < 0.01$; and *** → $p < 0.001$.

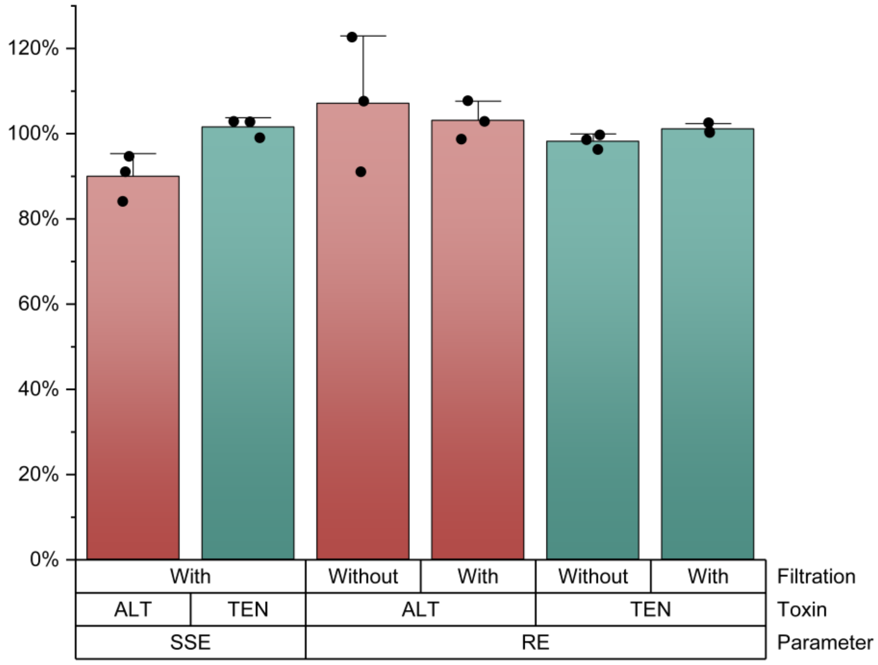


**Figure S2:** Signal suppression and enhancement (SSE) and extraction recovery with and without filtration (RE) for ALT and TEN in the cell culture medium of primary hepatocytes. Each bar represents the mean value of three technical replicates, and the error bar depicts the respective standard deviation.

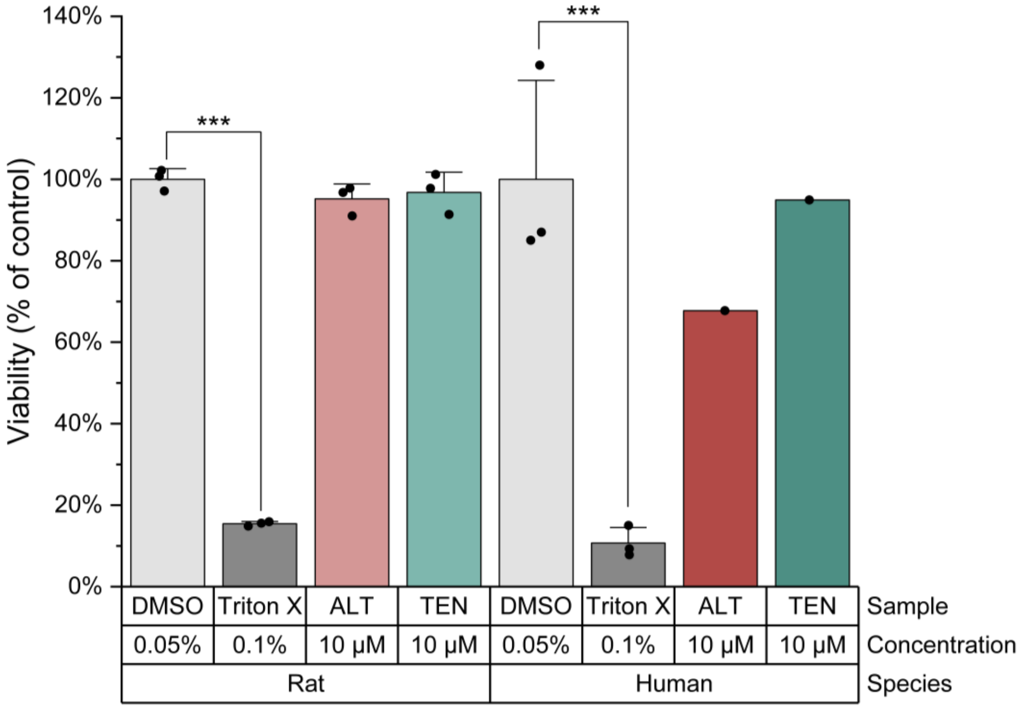


**Figure S3:** Viability of rat and human primary hepatocytes at the end of the kinetic incubated period (4 h) for ALT and TEN. Triton X-100 (0.1%) was used as positive control showing a significant decrease in viability (around ca. 80-90% decrease). Each bar represents the mean value and standard deviation of three replicates which are also depicted. For ALT and TEN in primary human hepatocytes, only one replicate was conducted, therefore, no statistical analysis could be carried out for these conditions. After testing for normality according to Shapiro-Wilk, one-way ANOVA, followed by Fisher's LSD post-hoc test was performed. The significance levels are marked as follows: * → $0.01 < p < 0.05$; ** → $0.001 < p < 0.01$; and *** → $p < 0.001$.

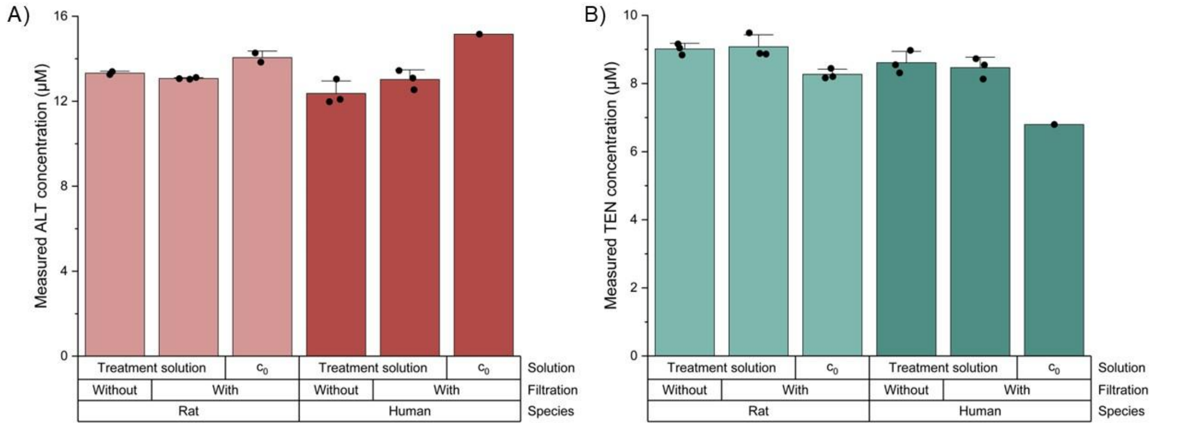


**Figure S4:** ALT (A) and TEN (B) concentrations measured in the treatment solutions by the targeted LC-MS/MS method used for incubation with primary rat and human hepatocytes, including the initial concentration ($c_0$). Each bar represents the mean value of three technical replicates from three biological replicates, with error bars indicating the standard deviation among biological replicates. For human hepatocytes, $c_0$ was determined from a single biological replicate. Outliers were identified and excluded using Nalimov's test. After testing for normality according to Shapiro-Wilk, one-way ANOVA, followed by Fisher's LSD post-hoc test was performed. No significant differences were found between the same samples with and without syringe filtration.

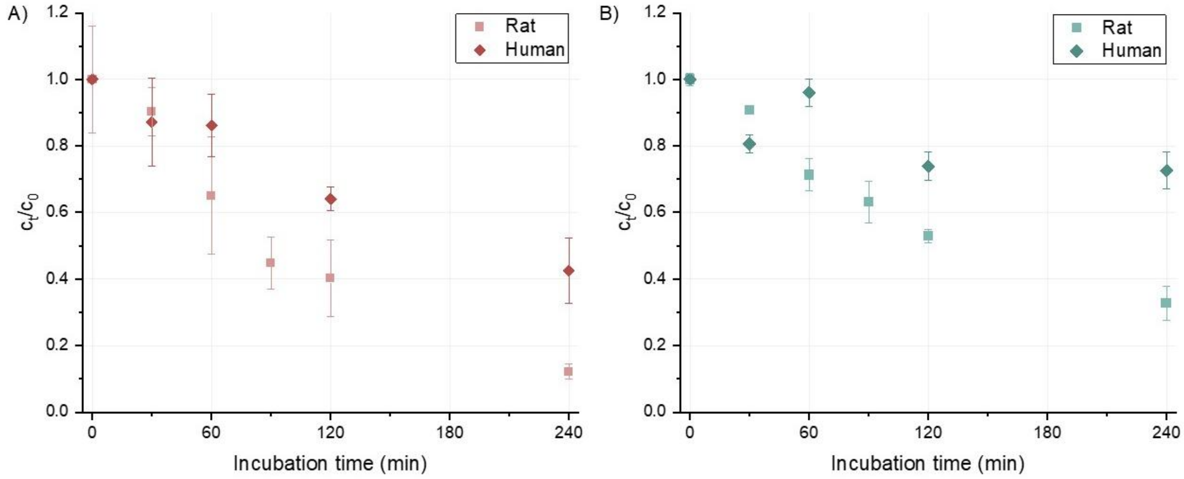


**Figure S5:** ALT (A) and TEN (B) concentrations measured in primary rat and human hepatocytes over 0–4 hours of incubation ($c_t$), normalized to the initial concentration ($c_0$). Each data point represents the mean of three technical replicates from three biological replicates, with error bars indicating the standard deviation among biological replicates. For human hepatocytes, $c_0$ was determined from a single biological replicate. Outliers were identified and excluded using Nalimov's test.

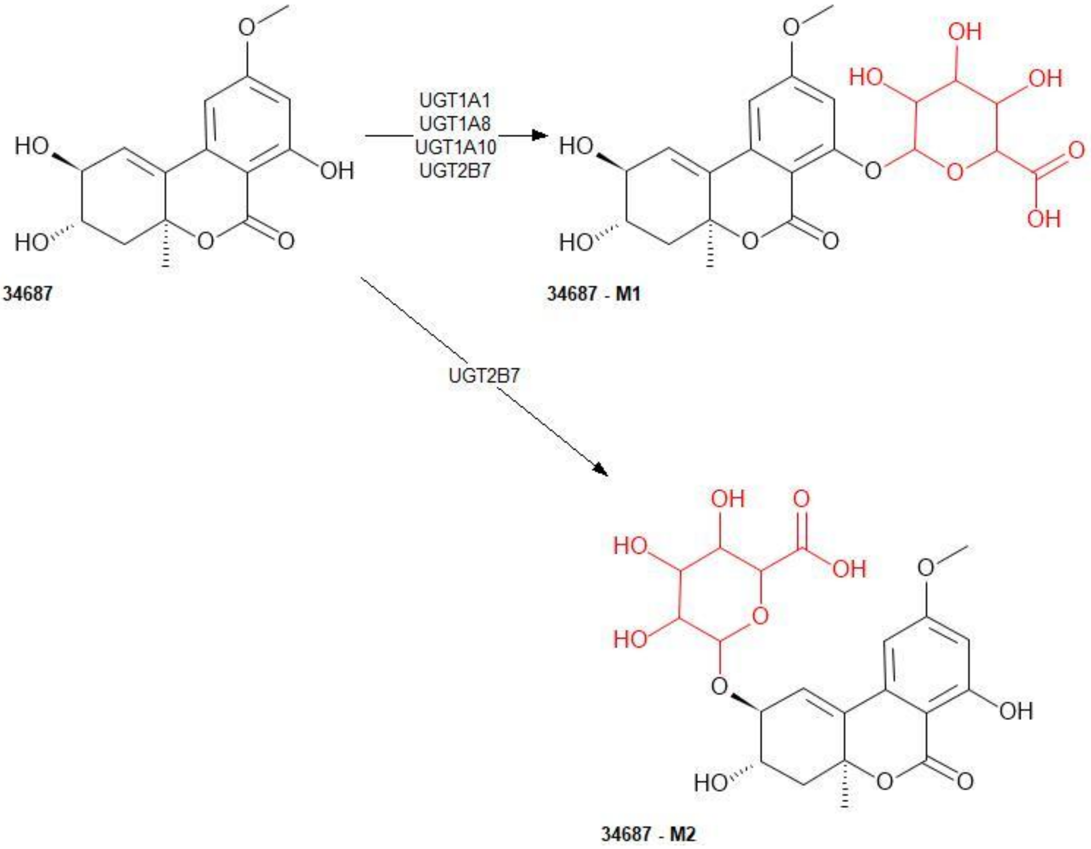


**Figure S6:** *In silico* predicted metabolite pathway for ALT

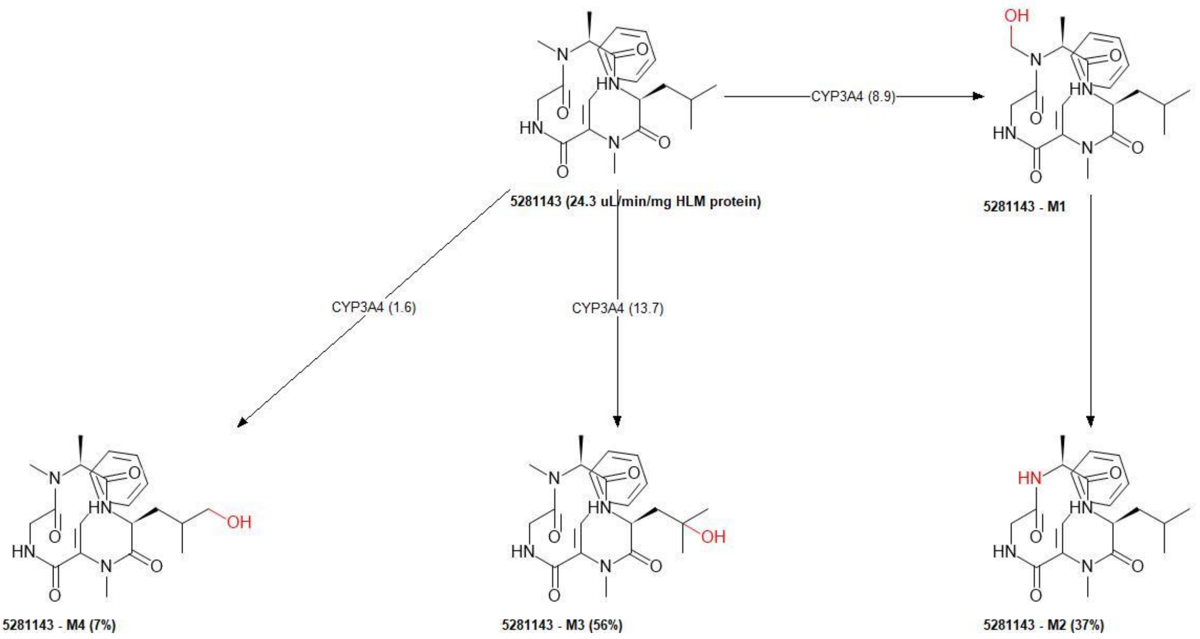


**Figure S7:** *In silico* predicted metabolite pathway for TEN

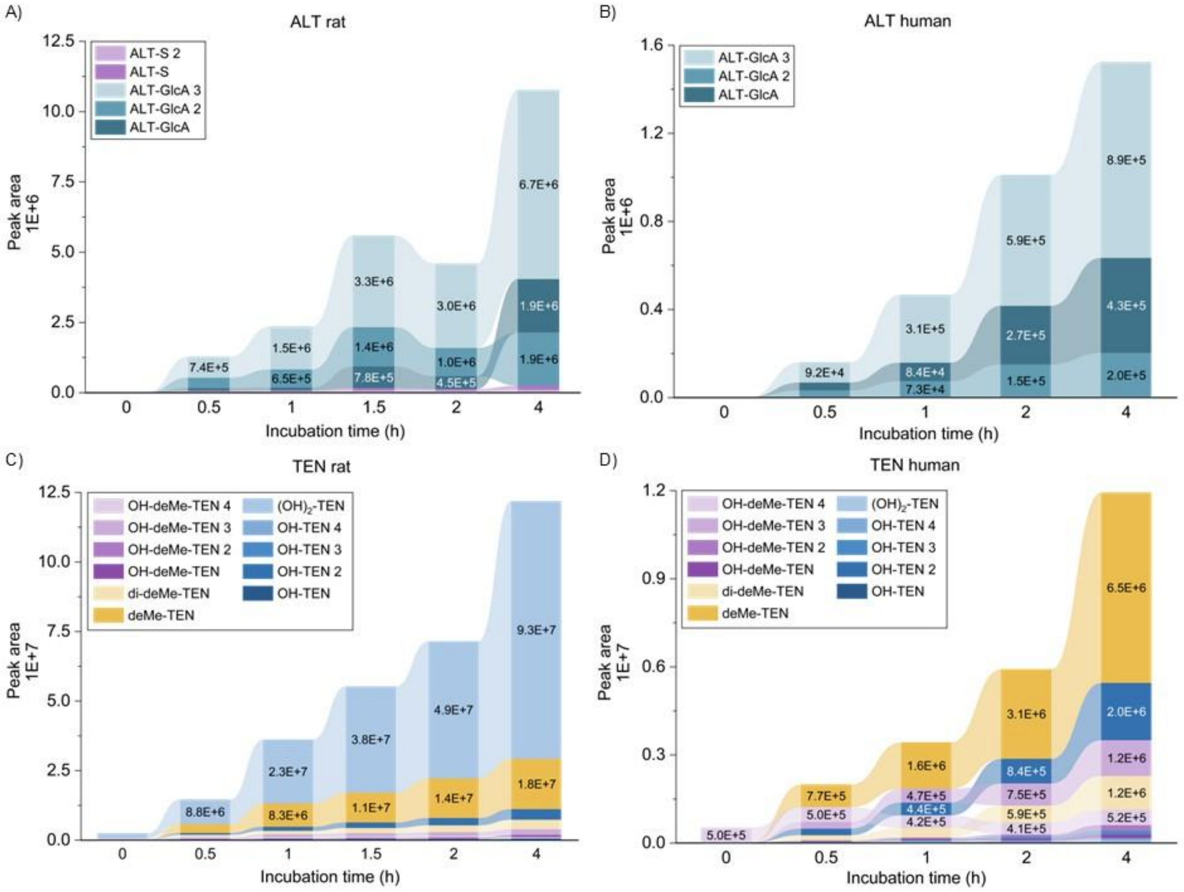


**Figure S8:** Interspecies differences in the metabolite profile in primary rat and human hepatocytes occurring after up to 4 h of incubation with 10 µM ALT (A-B) or TEN (C-D) measured via the targeted LC-MS/MS method. Each section shows the average of three independent experiments and metabolites are depicted in the order of their peak area. Please note the differences in the y-axes.

**Table S1:** Inclusion list for ALT and TEN in positive and negative mode

| **Mass (*m/z*)** | **Formula (M)** | **Formula type** | **Species** | **CS (z)** | **Polarity** | **Start (min)** | **End (min)** | **Comment** |
|---|---|---|---|---|---|---|---|---|
| 293.10196 | $C_{15}H_{16}O_6$ | Chemical formula | $[M+H]^+$ | 1 | Positive | 5.10 | 5.70 | ALT-pos |
| 415.23398 | $C_{22}H_{30}N_4O_4$ | Chemical formula | $[M+H]^+$ | 1 | Positive | 5.80 | 6.30 | TEN-pos |
| 291.08741 | $C_{15}H_{16}O_6$ | Chemical formula | $[M-H]^-$ | 1 | Negative | 2.00 | 26.00 | ALT-neg |
| 413.21943 | $C_{22}H_{30}N_4O_4$ | Chemical formula | $[M-H]^-$ | 1 | Negative | 2.00 | 26.00 | TEN-neg |

**Table S2:** Mass spectrometric parameters of the targeted LC-MS/MS measurements of ALT, TEN, and their metabolites. Transitions highlighted in bold were used for quantification/depiction of relative peak areas. Molecules without highlighted transitions were incorporated in the analytical method but were not detected. Abbreviations: ALT: altenuene; OH-ALT: hydroxy-ALT; ALT-GlcA: ALT-glucuronide; ALT-S: ALT-sulfate; TEN: tentoxin; OH-TEN: hydroxy-TEN; TEN-S: TEN-sulfate; TEN-GlcA: TEN-glucuronide; deMe-TEN: demethylated TEN; di-deMe-TEN: TEN demethylated in two positions; $(OH)_2$-TEN: dihydroxy-TEN; OH-deMe-TEN: hydroxylated demethylated TEN; DP: declustering potential; EP: entrance potential; CE: collision energy; CXP: cell exit potential.

| Molecule name | Q1 mass | Q3 mass | Dwell time (msec) | DP (V) | EP (V) | CE (V) | CXP (V) |
|---|---|---|---|---|---|---|---|
| ALT | **291.1** | **203.0** | **10** | **-110** | **-10** | **-45** | **-10** |
| | 291.1 | 247.1 | 10 | -110 | -10 | -30 | -10 |
| | 291.1 | 229.0 | 10 | -110 | -10 | -35 | -10 |
| | 291.1 | 161.0 | 10 | -110 | -10 | -55 | -10 |
| OH-ALT | 307.1 | 291.1 | 10 | -100 | -10 | -20 | -10 |
| | 307.1 | 276.0 | 10 | -100 | -10 | -30 | -10 |
| | 307.1 | 203.1 | 10 | -100 | -10 | -44 | -10 |
| ALT-GlcA | **467.1** | **291.0** | **10** | **-70** | **-10** | **-30** | **-10** |
| | 467.1 | 276.0 | 10 | -70 | -10 | -55 | -10 |
| | 467.1 | 203.0 | 10 | -70 | -10 | -80 | -10 |
| ALT-S | **371.0** | **291.1** | **10** | **-70** | **-10** | **-20** | **-10** |
| | 371.0 | 276.0 | 10 | -70 | -10 | -30 | -10 |
| TEN | **413.1** | **271.0** | **10** | **-70** | **-10** | **-22** | **-10** |
| | 413.1 | 141.0 | 10 | -70 | -10 | -26 | -10 |
| OH-TEN | **429.2** | **287.3** | **10** | **-70** | **-10** | **-25** | **-10** |
| | 429.2 | 141.0 | 10 | -70 | -10 | -30 | -10 |
| | 429.2 | 305.2 | 10 | -70 | -10 | -25 | -10 |
| TEN-S | 493.0 | 413.1 | 10 | -100 | -10 | -20 | -10 |
| | 493.0 | 141.0 | 10 | -100 | -10 | -26 | -10 |
| | 493.0 | 271.0 | 10 | -100 | -10 | -22 | -10 |
| TEN-GlcA | 589.2 | 413.0 | 10 | -100 | -10 | -20 | -10 |
| | 589.2 | 141.0 | 10 | -100 | -10 | -26 | -10 |
| | 589.2 | 271.0 | 10 | -100 | -10 | -22 | -10 |
| deMe-TEN | **399.2** | **127.0** | **10** | **-70** | **-10** | **-30** | **-10** |
| | 399.2 | 271.0 | 10 | -70 | -10 | -20 | -10 |
| | 399.2 | 215.3 | 10 | -70 | -10 | -30 | -10 |
| di-deMe-TEN | **385.2** | **218.2** | **10** | **-70** | **-10** | **-20** | **-10** |
| | 385.2 | 201.2 | 10 | -70 | -10 | -30 | -10 |
| | 385.2 | 367.2 | 10 | -70 | -10 | -20 | -10 |
| | 385.2 | 257.2 | 10 | -70 | -10 | -30 | -10 |
| | 385.2 | 342.3 | 10 | -70 | -10 | -30 | -10 |
| $(OH)_2$-TEN | **445.2** | **415.3** | **10** | **-70** | **-10** | **-50** | **-10** |
| | 445.2 | 385.3 | 10 | -70 | -10 | -50 | -10 |
| | 445.2 | 127.0 | 10 | -70 | -10 | -40 | -10 |
| | 445.2 | 271.0 | 10 | -70 | -10 | -30 | -10 |
| OH-deMe-TEN | **415.2** | **127.0** | **10** | **-70** | **-10** | **-30** | **-10** |
| | 415.2 | 287.0 | 10 | -70 | -10 | -20 | -10 |
| | 415.2 | 215.3 | 10 | -70 | -10 | -30 | -10 |
| | 415.2 | 397.2 | 10 | -70 | -10 | -20 | -10 |
| | 415.2 | 355.2 | 10 | -70 | -10 | -20 | -10 |

**Table S3:** *In silico* molecular weight of the putative phase I and II biotransformation products of ALT and TEN.

| Analytes | Proposed structure | Identifier | Molecular weight prediction | Prediction recovery (%) |
|---|---|---|---|---|
| ALT | 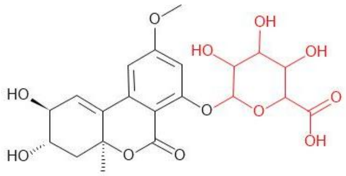 | M1<br>ALT-GlcA<br>altenuene-glucuronide | 468.417 | / |
| | 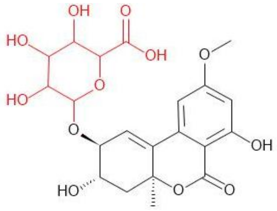 | M2<br>ALT-GlcA<br>altenuene-glucuronide | 468.417 | / |
| TEN | 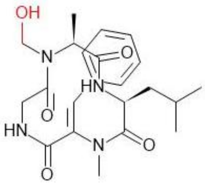 | M1<br>OH-TEN<br>hydroxy-tentoxin | 430.2209 | / |
| | 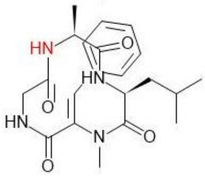 | M2<br>deMe-TEN<br>demethylated tentoxin | 400.2104 | 37 |
| | 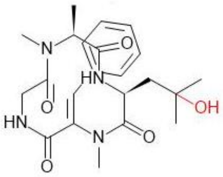 | M3<br>OH-TEN<br>hydroxy-tentoxin | 430.2209 | 7 |
| | 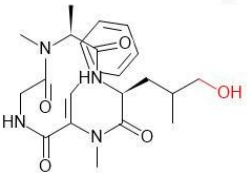 | M4<br>OH-TEN<br>hydroxy-tentoxin | 430.2209 | 56 |

**Table S4:** Postulated metabolic transformation of ALT formed in primary rat hepatocytes (PRH), retention time (RT), elemental composition, accurate mass of molecular ion (hydrogen adduct in positive-ionization mode $[M + H]^+$, hydrogen loss in negative-ionization mode $[M-H]^-$), deviation from theoretical accurate mass, using UHPLC-HRMS. No DDA – no data-dependent acquisition, no fragmentation triggered due to low intensity.

| ID | Proposed Biotransformation | Formula | $[M+H]^+$ | | | | $[M-H]^-$ | | | |
|---|---|---|---|---|---|---|---|---|---|---|
| | | | **RT (min)** | **Measured *m*/*z*** | **Mass Error (ppm)** | **Diagnostic Ions (*m/z*)** | **RT (min)** | **Measured *m*/*z*** | **Mass Error (ppm)** | **Diagnostic Ions (*m/z*)** |
| **M0** | ALT altenuene | $C_{15}H_{16}O_6$ | 12.2 | 293.1021 | 0.48 | 293.1016; 275.0914; 257.0808; 229.0861; 201.0912; 69.0343 | 12.3 | 291.0879 | 1.72 | 276.0637; 248.0690; 203.0348; 161.0240; 67.2128 |
| **M1** | ALT-S altenuene-sulfate | $C_{15}H_{16}O_9S$ | 11.5 | 373.0594 | 1.61 | No DDA | 11.5 | 371.0451 | 2.42 | 291.0872; 156.9959; 96.9589; 93.0335; 79.9561; 67.2134 |
| | | | 11.8 | 373.0595 | 1.88 | 257.0807; 239.0710; 227.0704; 214.0468; 99.0448; 67.2097; 56.9659 | 11.8 | 371.0452 | 2.69 | 291.0866; 255.0665; 156.9957; 96.9590; 93.0336; 79.9562; 67.2118 |

**Table S5:** continued

| ID | Proposed Biotransformation | Formula | $[M+H]^+$ | | | | $[M-H]^-$ | | | |
|---|---|---|---|---|---|---|---|---|---|---|
| | | | RT (min) | Measured *m/z* | Mass Error (ppm) | Diagnostic Ions (*m/z*) | RT (min) | Measured *m/z* | Mass Error (ppm) | Diagnostic Ions (*m/z*) |
| **M2** | ALT-GlcA altenuene-glucuronide | $C_{21}H_{24}O_{12}$ | 8.1 | 469.1354 | 2.98 | 293.1022; 275.0915; 257.0801; 67.2110 | 8.1 | 467.1206 | 2.35 | 291.0875; 255.3057; 175.0250; 113.0232; 85.0284; 67.2116 |
| | | | 8.3 | 469.1352 | 2.56 | No DDA | 8.4 | 467.1205 | 2.14 | No DDA |
| | | | 10.7 | 469.1353 | 2.77 | 293.1053; 275.0910; 257.0795; 67.2110 | 10.8 | 467.1205 | 2.14 | 291.0872; 273.0772 ; 113.0233; 85.0279; 67.2123 |
| | | | 11.4 | 469.1348 | 1.70 | 293.1021; 275.0913; 257.0807; 67.2100 | 11.4 | 467.1203 | 1.71 | 291.0879; 273.0774; 175.0247; 113.0234; 85.0283; 67.2133 |

**Table S5:** Postulated metabolic transformation of ALT formed in primary human hepatocytes (PHH), retention time (RT), elemental composition, accurate mass of molecular ion (hydrogen adduct in positive-ionization mode $[M + H]^+$, hydrogen loss in negative-ionization mode $[M-H]^-$), deviation from theoretical accurate mass, using UHPLC-HRMS. No DDA – no data-dependent acquisition, no fragmentation triggered due to low intensity.

| ID | Proposed Biotransformation | Formula | $[M+H]^+$ | | | | $[M-H]^-$ | | | |
|---|---|---|---|---|---|---|---|---|---|---|
| | | | RT (min) | Measured *m/z* | Mass Error (ppm) | Diagnostic Ions (*m/z*) | RT (min) | Measured *m/z* | Mass Error (ppm) | Diagnostic Ions (*m/z*) |
| **M0** | ALT altenuene | $C_{15}H_{16}O_6$ | 12.3 | 293.1021 | 1.02 | 293.1361; 257.0806; 229.0858; 201.0908; 67.2084 | 12.3 | 291.0875 | 0.34 | 291.0874; 248.0687; 203.0342; 67.2098 |
| **M1** | ALT-S altenuene-sulfate | $C_{15}H_{16}O_9S$ | 11.6 | 373.0588 | 1.87 | No DDA | 11.6 | 371.0456 | 3.77 | No DDA |
| | | | 11.8 | 373.0598 | 2.68 | No DDA | 11.8 | 371.0451 | 2.42 | No DDA |
| **M2** | ALT-GlcA altenuene-glucuronide | $C_{21}H_{24}O_{12}$ | 8.1 | 469.1345 | 1.06 | 293.1022; 275. 0915; 257.0801; 67.2110 | 7.8 | 467.1197 | 0.43 | 291.0876; 241.6321; 165.9089; 113.0232; 67.2096 |
| | | | 8.3 | 469.1348 | 1.70 | No DDA | 8.2 | 467.1196 | 0.21 | No DDA |
| | | | 11.0 | 469.1349 | 1.92 | 293.1053; 275.0910; 257.0795; 67.2110 | 10.8 | 467.1198 | 0.64 | 291.0872; 229.0863; 67.2103 |
| | | | 11.5 | 469.1342 | 0.43 | 293.1021; 275.0913; 257.0807; 67.2100 | 11.4 | 467.1196 | 0.21 | 291.0865; 273.0759; 229.0872; 67.2118 |

**Table S6:** Postulated metabolic transformation of TEN formed in primary rat hepatocytes (PRH), retention time (RT), elemental composition, accurate mass of molecular ion (hydrogen adduct in positive-ionization mode $[M + H]^+$, hydrogen loss in negative-ionization mode $[M-H]^-$), deviation from theoretical accurate mass, using UHPLC-HRMS. No DDA – no data-dependent acquisition, no fragmentation triggered due to low intensity.

| ID | Proposed Biotransformation | Formula | $[M+H]^+$ | | | | $[M-H]^-$ | | | |
|---|---|---|---|---|---|---|---|---|---|---|
| | | | RT (min) | Measured *m/z* | Mass Error (ppm) | Diagnostic Ions (*m/z*) | RT (min) | Measured *m/z* | Mass Error (ppm) | Diagnostic Ions (*m/z*) |
| **M0** | TEN<br>tentoxin | $C_{22}H_{30}N_4O_4$ | 14.0 | 415.2346 | 1.44 | 312.1708; 302.1501; 256.1810; 217.0974; 171.1494; 132.0810; 86.0971; 58.0661 | 14.0 | 413.2204 | 2.42 | 369.2296; 271.1456; 214.0747; 141.0662; 113.0348 |
| **M1** | OH-TEN<br>hydroxy-tentoxin | $C_{22}H_{30}N_4O_5$ | 11.0 | 431.2296 | 1.62 | 302.1506; 285.1243; 189.1023; 132.0810; 58.0661 | 11.0 | 429.2152 | 2.1 | 287.1404; 215.0826; 141.0660; 109.0398 |
| | | | 11.7 | 431.2299 | 2.32 | 272.1655; 233.0925; 205.0970; 86.0973; 58.0661 | 11.6 | 429.2152 | 2.1 | 287.1409; 141.0661 |
| | | | 12.8 | 431.2299 | 2.32 | 233.0918; 205.0973; 86.0973; 58.0661 | 12.9 | 429.2151 | 1.9 | No DDA |
| | | | 13.4 | Trace | | | 13.4 | 429.2155 | 2.8 | No DDA |

**Table S6**: continued

| ID | Proposed Biotransformation | Formula | $[M+H]^+$ | | | | $[M-H]^-$ | | | |
|---|---|---|---|---|---|---|---|---|---|---|
| | | | **RT (min)** | **Measured *m/z*** | **Mass Error (ppm)** | **Diagnostic Ions (*m/z*)** | **RT (min)** | **Measured *m/z*** | **Mass Error (ppm)** | **Diagnostic Ions (*m/z*)** |
| **M2** | deMe-TEN demethylated tentoxin | $C_{21}H_{28}N_4O_4$ | 13.1 | 401.2188 | 1.25 | 217.0975; 189.1025; 132.0810; 86.0972; 58.0662 | 13.2 | 399.2046 | 2.0 | 381.1939; 271.1457; 232.1092; 127.0503; 99.0189 |
| **M3** | di-deMe-TEN di-demethylated tentoxin | $C_{20}H_{26}N_4O_4$ | 12.1 | 387.2032 | 1.29 | No DDA | 12.2 | 385.1890 | 2.33 | 294.1339; 218.0933; 201.0665; 127.0503; 99.0189 |
| **M4** | $(OH)_2$-TEN double hydroxylated tentoxin | $C_{22}H_{30}N_4O_6$ | 11.9 | 447.2248 | 2.23 | 235.1079; 132.0810; 86.0972 | 13.2 | 445.2102 | 2.02 | 399.2037; 271.1452; 127.0503 |
| **M5** | OH-deMe-TEN hydroxylated demethylated tentoxin | $C_{21}H_{28}N_4O_5$ | 10.6 | 417.2140 | 1.92 | 217.0974; 189.1024; 132.0810 | 10.6 | 415.1996 | 2.17 | 287.1403; 199.1083; 127.0504; 99.0189 |

**Table S7:** Postulated metabolic transformation of TEN formed in primary human hepatocytes (PHH), retention time (RT), elemental composition, accurate mass of molecular ion (hydrogen adduct in positive-ionization mode $[M + H]^+$, hydrogen loss in negative-ionization mode $[M\text{-}H]^-$), deviation from theoretical accurate mass, using UHPLC-HRMS. No DDA – no data-dependent acquisition, no fragmentation triggered due to low intensity.

| ID | Proposed Biotransformation | Formula | $[M+H]^+$ | | | | $[M\text{-}H]^-$ | | | |
|---|---|---|---|---|---|---|---|---|---|---|
| | | | **RT (min)** | **Measured *m/z*** | **Mass Error (ppm)** | **Diagnostic Ions (*m/z*)** | **RT (min)** | **Measured *m/z*** | **Mass Error (ppm)** | **Diagnostic Ions (*m/z*)** |
| **M0** | TEN tentoxin | $C_{22}H_{30}N_4O_4$ | 14.1 | 415.2342 | 0.48 | 312.1706; 302.1497; 217.0973; 199.1441; 189.1022; 132.0809; 86.0971; 58.0660 | 14.1 | 413.2195 | 0.24 | 271.1448; 214.0733; 141.0659 |
| **M1** | OH-TEN hydroxy-tentoxin | $C_{22}H_{30}N_4O_5$ | 11.1 | 431.2297 | 1.85 | 302.1490; 217.0969; 189.1022; 132.0809; 67.2081; 58.0661 | 11.1 | 429.2145 | 0.47 | 287.1404; 141.0659 |
| | | | 11.8 | 431.2296 | 2.09 | 67.2102; 58.0661 | 11.8 | 429.2144 | 0.23 | 287.1406; 141.0657; 113.0346 |
| | | | 12.5 | 431.2299 | 2.32 | No DDA | 13.0 | 429.2146 | 0.70 | No DDA |
| | | | 13.0 | 431.2292 | 0.70 | No DDA | 13.5 | 429.2147 | 0.93 | No DDA |
| **M2** | deMe-TEN demethylated tentoxin | $C_{21}H_{28}N_4O_4$ | 13.3 | 401.2190 | 1.74 | 217.0977; 189.1028; 132.0810; 86.0973; 56.0661 | 13.3 | 399.2040 | 0.50 | No DDA |

**Table S7:** continued

| ID | Proposed Biotransformation | Formula | $[M+H]^+$ | | | | $[M-H]^-$ | | | |
|---|---|---|---|---|---|---|---|---|---|---|
| | | | **RT (min)** | **Measured *m*/*z*** | **Mass Error (ppm)** | **Diagnostic Ions (*m/z*)** | **RT (min)** | **Measured *m*/*z*** | **Mass Error (ppm)** | **Diagnostic Ions (*m/z*)** |
| **M3** | di-deMe-TEN di-demethylated tentoxin | $C_{20}H_{26}N_4O_4$ | 12.3 | 387.2033 | 1.55 | 132.0810; 86.0972 | 12.3 | 385.1883 | 0.52 | No DDA |
| **M4** | $(OH)_2$-TEN double hydroxylated tentoxin | $C_{22}H_{30}N_4O_6$ | 12.0 | 447.2248 | 2.23 | 132.0810; 86.0972; 67.2093 | 12.1 | 445.2098 | 1.35 | 229.1204 |
| | | | | | | | 13.3 | 445.2095 | 0.67 | 399.2009; 74.0058 |
| **M5** | OH-deMe-TEN hydroxylated demethylated tentoxin | $C_{21}H_{28}N_4O_5$ | 10.8 | 417.2139 | 1.68 | No DDA | 10.7 | 415.1990 | 0.72 | 287.1391; 127.0499 |
